# Disentangling Multidimensional Spatio-Temporal Data into their Common and Aberrant Responses

**DOI:** 10.1101/004259

**Authors:** Young Hwan Chang, Jim Korkola, Dhara N. Amin, Mark Moasser, Jose M. Carmena, Joe W. Gray, Claire J. Tomlin

## Abstract

With the advent of high-throughput measurement techniques, scientists and engineers are starting to grapple with massive data sets and encountering challenges with how to organize, process and extract information into meaningful structures. Multidimensional spatio-temporal biological data sets such as time series gene expression with various perturbations over different cell lines, or neural spike trains across many experimental trials, have the potential to acquire insight across multiple dimensions. For this potential to be realized, we need a suitable representation to understand the data. Since a wide range of experiments and the unknown complexity of the underlying system contribute to the heterogeneity of biological data, we propose a method based on Robust Principal Component Analysis (RPCA), which is well suited for extracting principal components when there are corrupted observations. The proposed method provides us a new representation of these data sets in terms of a common and aberrant response. This representation might help users to acquire a new insight from data.

**Author Summary:** One of the most exciting trends and important themes in science and engineering involves the use of high-throughput measurement data. With different dimensions, for example, various perturbations, different doses of drug or cell lines characteristics, such multidimensional data sets enable us to understand commonalities and differences across multiple dimensions. A general question is how to organize the observed data into meaningful structures and how to find an appropriate similarity measure. A natural way of viewing these complex high dimensional data sets is to examine and analyze the large-scale features and then to focus on the interesting details. With this notion, we propose an RPCA-based method which models common variations as approximately the low-rank component and anomalies as the sparse component. We show that the proposed method is able to find distinct subtypes and classify data sets in a robust way without any prior knowledge by separating these common responses and abnormal responses.

## Introduction

Over the last years, the use of high-throughput measurement data has become one of the most exciting trends and important themes in science and engineering. This is becoming increasingly important in biology. However, handling and analyzing biological data have challenges all of their own because the data sets are typically heterogeneous. Biological data can not only stem from a wide range of experiments such as inhibitions/stimulations, different doses of drugs, and various cell lines (Figure 1) but also represent the (unknown) complexity of the underlying system [1].

**Figure 1.**
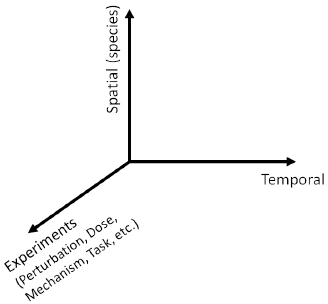
Multi-dimensional Spatiotemporal data where we consider various experiments with different perturbations, doses, mechanism, tasks, etc.

With the explosion of the amount of various biological data, a general question is how to organize the observed data into meaningful structures and how to find an appropriate similarity (or dissimilarity) measure which is critical to the analysis. Since such multidimensional data have the potential to provide insight across multiple dimensions, these data can enable users to start to develop models and draw hypotheses that not only describe the spatial and temporal dynamics of the biological system but also inform them about commonalities and differences across dimensions. A significant challenge for creating suitable representations is to continue handling large data sets and effectively deal with the growing diversity and quantity of the data set.

A natural way of viewing these complex high dimensional data sets is to examine and analyze the large-scale features and then to focus on the interesting details. The potential of clustering to reveal biologically meaningful patterns in microarray data was first realized and demonstrated in an early paper by Eisen *et al* [2]. Thereafter, in many biological applications, different methods have been used to analyze gene expression data and characterize gene functional behavior. Among various data-driven modeling approaches, clustering methods are widely used on these data to categorize genes with similar expression profiles. However, until recently, most studies have focused on the spatial, rather than temporal, structure of data. For instance, neural models are usually concerned with processing static spatial patterns of intensities without regard to temporal information [3]. Since many existing data-driven modeling approaches such as clustering, classification or inference using biological data focus on static data, they have limitations in analyzing multi-dimensional spatio-temporal data sets.

Recently, much research has focused on time series high-throughput data sets. These data sets have the advantage of being able to identify dynamic relationships between genes or neurons since the spatio-temporal pattern results from integration of regulatory signals or electrochemical signals through the network over time. For example, time series gene-knockout experiment data sets provide the distinct possibility of observing the cellular mechanisms in action [4]. Also, these data sets help us to unravel the mechanistic drivers characterizing cellular response and to break down the genome into sets of genes involved in the related processes [5]. Moreover, instead of concentrating on steady state response, monitoring dynamic patterns provides a profoundly different type of information. For instance, several recent studies focus on the temporal complexity and heterogeneity of single-neuron activity in the premotor and motor cortices [3] [6] [7]. Moreover, since many current and emerging cancer treatments are designed to inhibit or stimulate a specific node (or gene) in the networks and alter signaling cascades, advancing our understanding of how the system dynamics of these networks is deregulated across cancer cells and finding subgroups of genes and conditions will ultimately lead to the more effective treatment strategies [8].

In this paper, we propose the Robust Principal Component Analysis (RPCA)-based method for analyzing spatio-temporal biological data sets. To demonstrate that our method helps users acquire insight efficiently and to emphasize that the proposed method can be applicable to various domains, we consider two different systems 1) neural population dynamics and 2) a gene regulatory network. Since the proposed method uses the common dynamic features in the spatio-temporal data set, it is important to arrange individual data sets in order to make them amenable to this analysis.

## Background

### Motivation

#### 1) Neural Population Dynamics

Neural ensemble activity is typically studied by averaging noisy spike trains across multiple experimental trials to obtain an approximate neural firing rate that varies smoothly over time. However, if neural activity is more a reflection of internal neural dynamics rather than response to external stimulus, the time series of neural activity may differ even when the subject is performing nominally identical tasks [7]. In [6], Churchland *et al*. showed that neural activity patterns in the primary motor cortex and dorsal premotor cortex of the macaque brain associated with nearly identical velocity profiles can be very different. This is particularly true of behavioral tasks involving perception, decision making, attention, or motor planning. In these settings, it is critical not to average the neural data across trials, but to analyze it on a trial-by-trial basis [3]. Moreover, stimulus representations in some sensory systems are characterized by the precise spike timing of a small number of neurons [10] [11] [12], suggesting that the details of operations in the brain are embedded not only in the overall neural spike rate, but also in the timings of spikes.

The motor and premotor cortices have been extensively studied but their dynamic response properties are poorly understood [3]. Moreover, the role of motor cortex in arm movement control is still unclear, with experimental evidence supporting both low-level muscle control as well as high-level kinematic parameters. We can define the motor cortical activity, which represents movement parameters as per equation (1), and the dynamical system that generates movements as per equation (2) [3]:

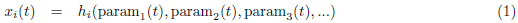

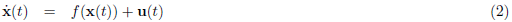

where *x*_*i*_(*t*) is the firing rate of neuron *i* at time *t*, *h*_*i*_ is its tuning function, and each param_*j*_ may represent a movement parameter such as hand velocity, target position or direction. In (2), **x** ∈ ℝ^*n*^ is a vector describing the firing rate of all neurons where *n* is the number of neurons, 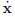 is its derivative, *f* is an unknown function, and **u** is an external input. In (2), neural activity is governed by the underlying dynamics *f*(·), so the characteristics of dynamical system should be present in the population activity. Also, if we align spatio-temporal neural activity as shown in Figure 2(b), we may extract these characteristics.

#### 2) Gene Regulatory Network

In microarray data, missing and corrupted data, including arbitrary corruptions by human error during biological experiments, are quite common, and not uniform across samples. Two strategies for dealing with missing values are either to modify clustering methods so that they can deal with missing values, or impute a “complete” data set before clustering [13].

Consider collections of time series gene expression of breast cancer cell lines or microarray data sets from pathway-targeted therapies involving gene knockout experiments. When a specific gene is perturbed as shown in Figure 2(c), the broad gene expression levels of other genes might be perturbed over time. Thus, comparing gene expression levels in the perturbed system with those in the unperturbed system reveals the extra information that is the different cellular mechanisms in action. A dynamical system of the gene regulatory network can be modelled as follows:

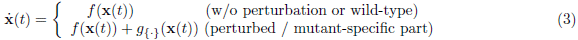

where 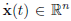 denotes the concentrations of the rate-limiting species, *x*̇(*t*) represents the change in concentration of the species over time *t*, *n* is the number of genes, *f*(·) represents the vector field of the typical dynamical system (or wild-type) and *g*_{·}_(·) represents an additional perturbation or mutant-specific vector field (blue and red edges in Figure 2(c)). In other words, we have a unified model for wild-type cell line, 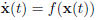 and in the mutant or perturbation case, we invoke a single change to the network topology or add a single influence for a specific gene. Here, additional vector fields such as *g*_LAP_(·), *g*_Akti_(·) and *g*_M_(·) are assumed to be sparse (i.e., affect only a single gene expression). Although these additional vector fields affect only a single gene expression at time *t*, their influence can be propagated through the network over time.

**Figure 2.**
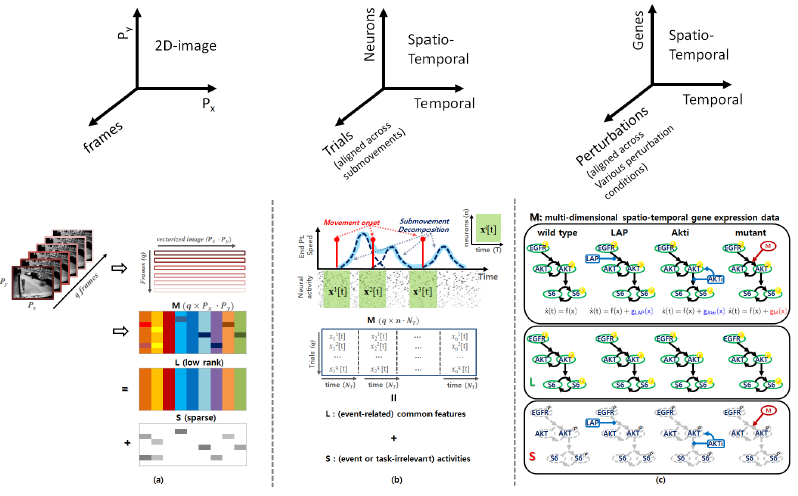
Conceptual representation: (a) RPCA applied to computer vision. A typical example of video surveillance where the low-rank component represents the unchanging background and the sparse component represents the movements in the foreground. (b) RPCA applied to neural systems. The low-rank component putatively represents (submovement relevant) neural signatures and the sparse component represents neural activity unrelated to submovement onset. (c) Collections of gene-knockout experiments and mutant-specific part representations (breast cancer signaling pathway) with `wild-type`, `Lapatinib` treatment, `Akt` inhibitor and `mutant` cell lines where solid black edges represent common network topology, and blue and red edges represent a single change of the network topology for perturbations or mutant cell lines.

### Robust Principal Component Analysis (RPCA)

In the computer vision literature [9], an interesting separation problem is introduced where the observed data matrix can be decomposed into an unseen low-rank component and an unseen sparse component. The method called Robust Principal Component Analysis (RPCA) is a provably correct and efficient algorithm for the recovery of low-dimensional linear structure from non-ideal observations, incorporating for example, occlusions, malicious tampering, and sensor failures.

In video surveillance, we need to identify activities that stand out from the background given a sequence of video frames [9]. Figure 2(a) shows that if we stack the video frames as rows of a matrix 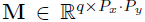 where *q* is the number of frames for a given time window, and *P*_*x*_ and *P*_*y*_ represent the number of pixels of 2-D images respectively, then across each row of **M**, there exists a common component that is the stationary background and a changing component which are the moving objects in the foreground at each image frame. Here, the data matrix **M** is an input for RPCA and the output is both the stationary background represented as a matrix **L** ∈ ℝ^*q* × *P_x_ · P_y_*^ and the moving objects in the foreground represented as a matrix **S** ∈ ℝ^*q* × *P_x_ · P_y_*^. With only one frame, the moving objects cannot be identified from the stationary background. However, by stacking all the vectorized frames such that all the frames align across the column direction as shown in Figure 2(a), we can identify the stationary backgrounds which are common variations, and then capture the moving objects which are sparse components for each frame.

With this notion, suppose we are given a large data matrix **M**, which has principal components in the low-rank component and may contain some anomalies in the sparse component. Mathematically, it is natural to model the common variations as approximately the low-rank component **L**, and the anomaly as the sparse component **S**. In [9], Candès *et al*. formulate this as follows:

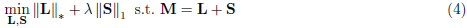

where ‖**L**‖_∗_ denotes the so-called nuclear norm of the matrix **L**, which is the sum of the singular value of **L**, and ‖**S**‖_1_ = ∑_*ij*_ |**S**_*ij*_| represents *l*_1_-norm of **S**. The turning parameter λ may be varied to put more importance on the rank of **L** or the sparseness of **S**. Choosing the tuning parameter λ to be 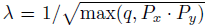 works well in practice [9]. However, appropriate choice of λ remains an open problem. Thus, we can use λ as a turning parameter to trade off more importance between **L** and **S**.

### How to Construct the Data Matrix M

In the video surveillance example shown in Figure 2(a), each row of **M** represents the vectorized 2-D images at each time frame. Since each image consists of the stationary background (**L**_*i*,:_) and the moving objects in the foreground (**S**_*i*,:_) at each time *i*, we denote **M** as follows:

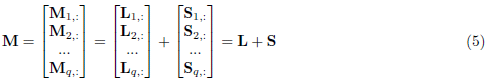

where **M**_*i*,:_ represents the *i*-th row of **M**. If there were no moving object in the foreground and no variation for a given video sequence (i.e., ∀*i*, **S**_*i*,:_ = 0), **L**_*i*,:_(= **L**_*j*,:_(*i* ≠ *j*)) would represent the common stationary background. On the other hand, if not (i.e., **S**_*i*,:_ ≠ 0), **M** represents the aligned corrupted measurements **M**_*i*,:_. Although the measurements are corrupted by moving objects in the foreground, we are able to separate **L** and **S** under certain conditions [9].

#### 1) Neural Population Dynamics

Recall equation (2) and consider an experiment involving a non-human primate subject instructed to make visually-guided planar reaches with its hand. During the experiment, hand position and velocity, as well as the discharge of neurons from primary motor cortex and dorsal premotor cortex were recorded. See recerence [14] for details on the data sets. All procedures were conducted in compliance with the National Institute of Health Guide for Care and Use of Laboratory Animals and were approved by the University of California, Berkeley Institutional Animal Care and Use Committee. Then, hand velocity data were decomposed into a sum of minimum-jerk basis functions where submovement representation is a type of motor primitive; for example, the hand speed profile as a function of time resulting from arm movements can be represented by a sum of bell-shaped functions as shown in Figure 2(b), each of which is called a submovement [14] and denoted as different trials. In Figure 2(b), each red bar denotes submovement onset, i.e., when the subject triggers submovement.

Suppose we align the spatio-temporal neural activity 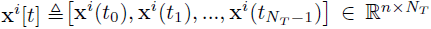 governed by (2) with submovement onset where the superscript *i* represents the *i*-th trial and N*_T_* represents the number of time points for the chosen time window. Then, M may be represented as follows:

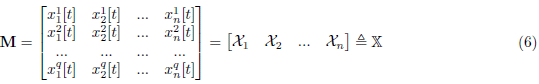

where 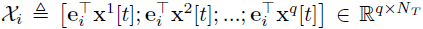 represents the temporal neural activity of the *i*-th neuron, **e**_*i*_ ∈ ℝ^*n*^ is a unit vector, and *q* is the trials or submovements. Thus, each row of M represents the vectorized vectorized spatio-temporal neural response for the each trial. Note that we align each spatio-temporal data set **x**^*j*^[*t*] with the same temporal condition (submovement onset) as shown in Figure 2(b) but we do not separate different types of submovement. For example, submovements with different reach directions, or with different ordinal positions in an overlapped series of submovements, are combined in our input matrix 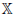. With the similar notion of the stationary background in video surveillance, some portion of the variability may reflect common dynamic features (**L**) corresponding to triggering submovement even though the responses of each neuron are corrupted by task-irrelevant neural responses (**S**) and may vary significantly across many trials.

#### 2) Gene Regulatory Network

Recall equation (3) and consider Figure 2(c). In (3), the vector field (*g*_{·}_) represents a single influence for a specific gene, yet this single influence can be propagated through the network over time. For example, when we inhibit *x*_*j*_, the gene expression levels of other genes can be perturbed over time. If *x*_*j*_ is connected with only few genes, this perturbation may only affect a small fraction of the total number of gene expression levels.

Similar to equation (6), we construct **M** using gene expression time series data with *q* different perturbations and/or different cell lines. Here, each row of 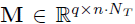 represents the vectorized time series gene expression 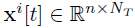 (*n*: the number of genes, *N*_*T*_: the number of time points and *q*: the number of different perturbation conditions including the number of different cell lines) and different rows represent spatio-temporal responses of different perturbations or different cell lines.

Since time series gene expression results from integration of regulatory signals constrained by the gene regulatory network, the input matrix **M** may reflect common dynamic response corresponding to the characteristics of the network structure. Intuitively, in video surveillance, if someone stays motionlessly in all the frames, the RPCA algorithm discriminates him as a low rank component. Unless he moves, we could not see the background because he always blocks the background. Similarly, in order to extract common response of gene regulatory network exactly, we should perturb the entire network arbitrarily and uniformly.

## Results

### Disentangling the Low-rank and Sparse components

In [9], Candès *et al*. discuss the identifiability issue. To make the problem meaningful, the low-rank component **L** must not be sparse. Another identifiability issue arises if the sparse matrix has low-rank. In many computer vision applications, practical low-rank and sparse separation gives visually appealing solutions.

However, for neural activity data, only a small subset of the whole ensemble of neurons is active at any moment as shown in Figure 3(left). Since **M** is sparse, the low-rank component might be sparse. Also, for the pathway targeted therapies, since gene regulatory networks are known to be sparse, a large subset of the whole ensemble of genes might be deactivated at any moment. Moreover, the original distributions of the amplitude of individual neuronal activities or gene expressions are highly skewed. For example, neural activities often form very eccentric clusters shown in Figure 3(left); some neurons are highly activated (30-40 spikes/sec) but others typically have only a few spikes per second. Similarly, gene expressions form very eccentric clusters since each gene expression shows different scales in practice.

**Figure 3.**
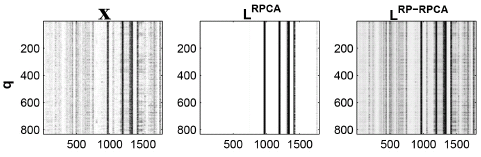
The low-rank matrices from both RPCA and RP-RPCA where 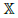 are input matrices and we choose *m* = *n* = 64 for the comparison (contrast represents activity of neuron. i.e., high contrast represents highly modulated neural activity and white color represents zero neural activity). (left) raw-data (center) low-rank component using RPCA and (right) low-rank component using RP-RPCA.

These imply that practical low-rank and sparse separation seems to be ambiguous and might present a challenge to achieving biologically meaningful solutions in both neural activity analyses and gene knockout experiment data sets. To remedy this identifiability issue, we propose the RPCA-based method in conjunction with Random Projection (RP); RP can de-sparsity the input data set and make a highly eccentric distribution more spherical so it makes the singular vectors of the low-rank matrix reasonably distributed. (see **Methods section: Random Projection (RP) and Identifiability** for details)

### Numerical Example

To illustrate the issue of identifiability and how RP can alleviate the issue, we consider a simple example: we generate a sparse low-rank input matrix 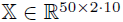 (*q* = 50, *n* = 2, *N*_*T*_ = 10) where the rank of 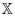 is 6 as shown in Figure S1(a). Note that in this example we chose the same dimension for the input 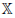 and 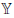 (refer to (7) and (8), no dimension reduction). This is done so that **Ψ** ∈ ℝ^*m*×*n*^ in equation (7) is invertible (we choose *m* = *n* and a nonsingular matrix **Ψ**), allowing us to compare the outputs of RPCA and RP-RPCA directly, as described below. Here, by using RP, we take advantage of de-sparsifying our input data and reducing the eccentric distribution. In general, choosing *m* < *n* makes 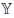 much denser because information is compressed by RP.

To evaluate the performance of separation into a low-rank and a sparse component, we add sparse corruption for 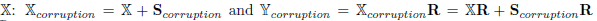 where 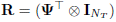 is the projection so 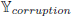 is the projected corrupted input 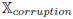. To compare the performance of RP-RPCA with RPCA, we first decompose 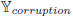 into its low-rank and sparse components. Then, we invert the projection:

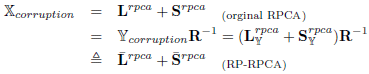

where we define 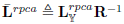 and 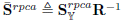.

Figure 4 shows statistics of both RPCA and RP-RPCA (in which RPCA is applied to the matrix 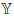) as a function of the tuning parameter λ in equation (4). In this example, 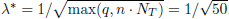. Since our input is still sparse in this example, the rank of both **L**^*rpca*^, 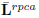 is 15 for 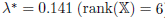. If we choose λ = 0.113 (discounting the penalty for sparse component), the ranks of **L**^*rpca*^, 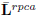 are approximately 6, which is the same as the rank of the original input 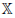. With this choice of λ, for RPCA we find that ‖**S**^*rpca*^‖ is much bigger than the original corruption signal 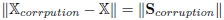. On the other hand, for RP-RPCA, we have 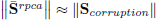. Therefore, for RP-RPCA, the separation of the low-rank component and sparse component is close to the true solution; for the original RPCA, there is mis-identification in both low-rank and sparse components *(more detailed information is provided in Figure S2)*.

**Figure 4.**
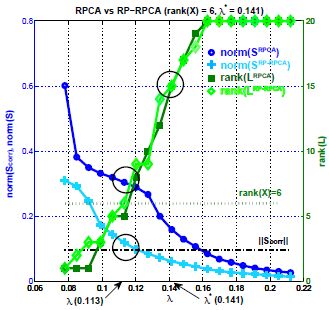
Statistics of a numerical example: we run RPCA for 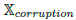 and 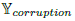 (We added sparse corruption to 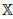). Left *y*-axis represents the norm of sparse component and the right *y*-axis shows the rank of **L** (more detailed information in Figure S1 and Figure S2).

### Application to Neural Data

Figure 3(left) shows the recorded neural activity aligned with submovement onset. The aligned neural activity shows that the ratios between units’ mean firing rates are fairly constant from the salient vertical striations in the plots and that temporal patterns exists across all the submovements. Also, as mentioned previously, the neural population activities are sparsely active (white color represents 0 spikes/sec) and show eccentric behavior; for example, some neurons have a much higher spiking rate than others.

Figure 3(middle)(right) show the low-rank matrix from both RPCA and RP-RPCA respectively (for simple comparison, we choose *m* = *n*). Since 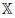 is sparse and has an eccentric distribution, the singular vectors may not be reasonably spread out. Applying RPCA directly to 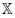 would result in the low-rank component being composed of only highly modulated neural activity (middle). On the other hand, RP-RPCA can extract the low-rank component from a more distributed set of neural dimensions than RPCA alone can. Also, the result of RP-RPCA gives a more visually appealing solution.

Since we extract neural features which represent common dynamic patterns across many experimental trials, we can use these features to detect and predict the onset of submovements. Here, we simply use the correlation between the extracted neural features and the neural signals. To accurately predict submovement onset times found by submovement decomposition, the correlation function should peak around the movement onset time. The following observations suggest the potential application of RP-RPCA to predict movement execution in a closed-loop Brain Machine Interface (BMI) system:

- **(observation 1)** Figure 5(a) represents the receiver operating characteristic (ROC) curve of the prediction of submovement onset time. We can see that the overall prediction performance based on RP-RPCA is better than the performance based on RPCA; we can reduce the false positive rate while increasing the true positive rate.
- **(observation 2)** Figure 5(b) shows the ROC curves of the prediction of submovement onset for different subjects or various tasks including center-out task and random-pursuit. This prediction could allow correction of movement execution errors in a closed-loop BMI system.

**Figure 5.**
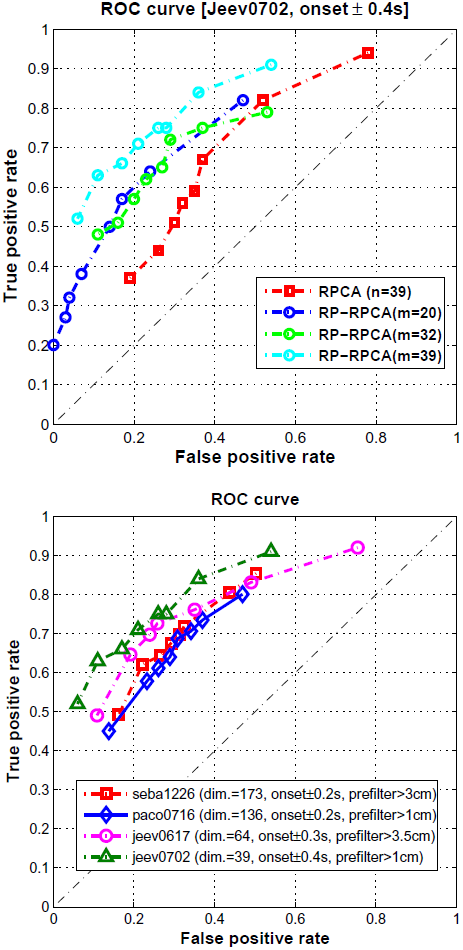
Receiver Operating Characteristic (ROC) curve of the prediction of submovement onset: (a) comparison between RPCA and RP-RPCA (target jumps task) (b) different monkeys or tasks where we prefiltered certain submovements with small amplitude in order to avoid artifacts of overfitting.

### Application to gene knockout experiments

To test the proposed RP-RPCA algorithm, we consider gene knockout experiments using `SKBR3` cell line [4] which has been used in studies of Human Epidermal Growth Factor Receptor2 (HER2) positive breast cancer. We chose this data set because it has 16 perturbations using a single cell line and contains 15 gene expressions with 4 time points as shown in Figure 6(top row). The middle row represents the low-rank component and the bottom row represents the highly aberrant sparse component. In raw data In raw data (top row), nearly all treatments show differential responses. However, the low-rank component (middle row) can be categorized into approximately 3-4 subtype responses, and the sparse component (bottom row) shows genomic aberration-specific responses.

**Figure 6.**
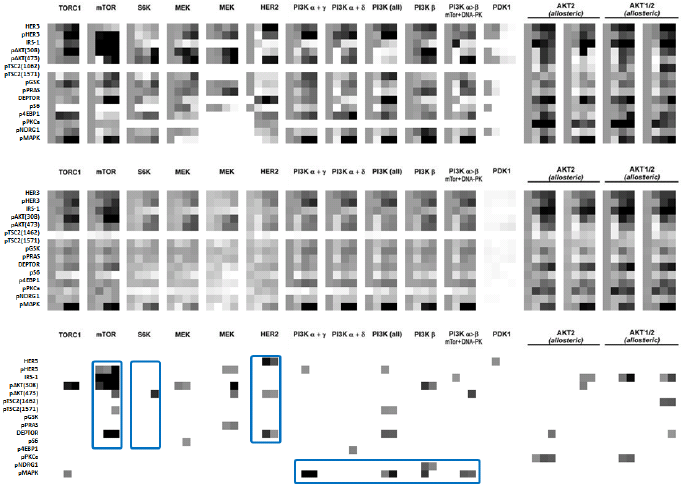
Gene knockout experiments [4](16 perturbations × 15 gene expressions × 4 time points [0, 1, 48, 72h]): (upper) raw data (middle) low-rank component and (lower) highly corrupted sparse component using threshold.

Also, the following observations suggest mechanisms of response and resistance which may inform unanticipated biological insight.

- (**observation 1**) `mTOR` inhibition (the second column in the bottom row) shows aberration responses in `DEPTOR, pHER3, IRS-1` and `pAkt(308, 473)`. In [15], `DEPTOR` is identified as an `mTOR`-interacting protein whose expression is negatively regulated by `mTORC1` and `mTORC2`; Also, Peterson *et al*. found that `DEPTOR` overexpression suppresses `S6K1` but it activates `Akt` by relieving feedback inhibition from `mTORC1` to `PI3K` signaling. Therefore, high `DEPTOR` expression is necessary to maintain `PI3K` and `Akt` activation and is consistent with the previous result [15].
- (**observation 2**) `HER2` inhibition (the sixth column in the bottom row) results in aberration responses of `HER3, pAkt(473)` and `DEPTOR`. Figure S3 represents an abstract model of `HER2` overexpressed breast cancer by biological interpretation. Since high `DEPTOR` expression represents low `mTORC1` and `mTORC2` [15], there are increasing activated `HER3` and `Akt` by relieving inhibition according to this model. The more interesting fact is that `PHLPP` is known to dephosphorylate `SER473` in `Akt` (i.e., partially inactivating the kinase) which is captured in the sparse component `pAkt`(473).
- (**observation 3**) `S6K` inhibition (the third column in the bottom row) results in aberration responses of `pAkt(473)`. Since `S6K` is located downstream of the `Akt-TSC2-mTORC` pathway, `S6K` inhibition captures only activation of `pAkt(473)`.
- (**observation 4**) `PI3K` inhibition (the 7th-11th columns in the bottom row) leads to increase phosphorylation of `MAPK`.

We separate the common response from the aberrant responses using the proposed method. Since abnormal behaviors or different responses to external stimuli or different cell lines can be extracted from the information available in the data set, we could cluster data correctly and reveal biological meaningful patterns (**see Methods section: Cluster Analysis** for details). Figure 7 shows the clustered result using existing hierarchical clustering (raw data **M**, *d*_*xy*_ in (9)) and the proposed method ([**L S**], *d*_*ϕψ*_ in (10)) respectively. We match the clustered results with graphical representation and our clustered result is more consistent with the known network structure and responses than the result of existing hierarchical
clustering.

**Figure 7.**
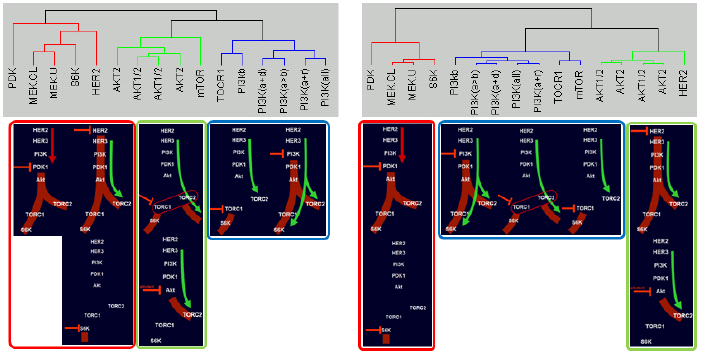
Clustered group: (left) hierarchical cluster and (right) the proposed method. Both clustered results compare with graphical representation generated by M. Moasser.

### Application to RPPA (Reverse Phase Protein Arrays) data set

Breast cancers are comprised of distinct subtypes which may respond differently to pathway-targeted therapies; collections of breast cancer cell lines show differential responses across cell lines and show subtype-, pathway-, and genomic aberration-specific responses [8]. These observations suggest mechanisms of response and resistance which differ across cell lines. Here, we use a data set generated in the Gray Lab using Reverse Phase Protein Arrays (RPPA) from the Mills Lab [16] which presents a time course analysis on 11 cell lines (all `HER2` amplified: `6 PI3K` mutant, `5 PI3K` wild-type) in response to `Lapatinib, Akt` inhibitor and combination of the two. The time course for RPPA is at 30min, 1h, 2h, 4h, 8h, 24h, 48h and 72h post-treatment.

As shown in Figure 8(top row), `Lapatinib` treatment results in down-regulation of a variety of phosphoproteins in the signaling pathway. From the raw data (**M**) or low-rank component (**L**), we can easily observe down-regulation and slow-recovery of the levels of activation, but the levels of activation are higher in the `PI3K` mutation cell lines. Treatment with `Akt` inhibitor leads to down-regulation of proteins (downstream of `Akt`) in all `HER2` amplified cell lines, although the amplitude of down-regulation is slightly less in cell lines with `PI3K` mutations. In the `PI3K` mutation cell lines, treatment with the combination of Lapatinib and Akt inhibitor leads to further down-regulation of the `Akt` signaling pathway but `Akt` levels are intermediate in comparison to those observed with inhibitor alone. Although these observations are still interesting, more interesting details might be in both the low-rank component **L** and the sparse component **S**:

- (**observation 1**) In the `PI3K` mutation with applying both inhibitors, full inhibition of `pS6RP` is observed and these results show the synergistic effect of `Lapatinib` and `Akt` inhibitor (in the bottom row, low-rank component).
- (**observation 2**) The main difference between wild-type and `PI3K` mutant is the response of `pS6RP` and `p70S6K`. For the wild-type cell lines, all treatments result in down-regulated pS6RP and p70S6K. However, for `PI3K` mutant cells, all treatments result in up-regulation `pS6RP` and `p70S6K` in the short-term (in the sparse component, red) and down-regulation in the long-term. Suppressing pS6RP relieves feedback inhibition and activates `Akt`. This difference makes `PI3K` mutation cells more resistant to `HER2` inhibitors than their wild-type counterparts. This finding is not obvious when we take a look at the raw data. Furthermore, our method makes our finding more convincing not by visually searching M, but by finding these effect automatically by separating common response (**L**) and aberrant behavior (**S**) by solving (4).
- (**observation 3**) `BT474` shows aberrant behavior as shown in Figure S4. This mutation has been reported to confer weak oncogenicity, unlike the other `PI3K` mutations.

**Figure 8.**
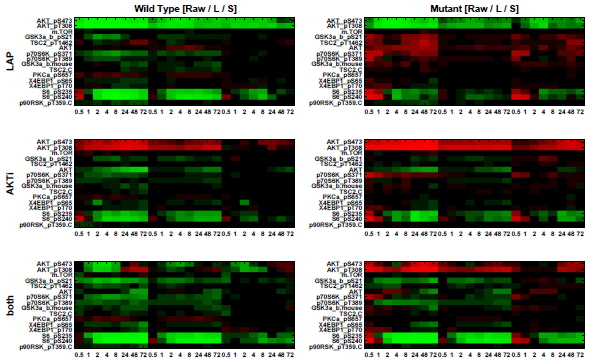
Heat maps showing average response based on both raw data and disentanglement result within subtype to targeted therapeutics: (1_*st*_ column) `HER2+/PI3K wild type`, (2_*nd*_ column) `HER2+/PI3K mutant`. Each column consists of average responses of raw RPPA, low-rank component and sparse component. Each row represents targeted therapeutics alone and in combination `(LAP, AKTi, both)`. In the PI3K mutation, we can see up-regulation of `S6 pS235, pS240` and `p70S6K pS371` in the short-term (in the sparse component, red) (*more detailed information for each cell line in Figure S4*).

Figure 9 shows the clustered result using existing hierarchical clustering and the proposed method respectively. Our clustered result is more robust and unaffected by different treatments due to the separation of the common responses. On the other hand, the clustered group based on existing hierarchical clustering changes across different treatments even though the characteristics of cell lines are not changing.

**Figure 9.**
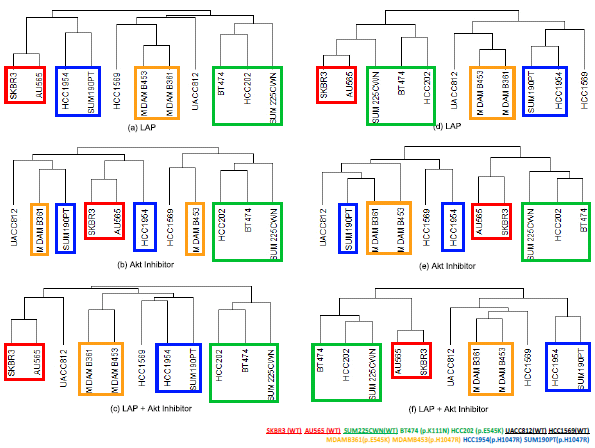
Clustered group using `RPPA` data set: (a,b,c) hierarchical cluster and (d,e,f) the proposed method.

## Discussion

Clustering and network inference are usually developed independently. For instance, until recently, most studies of gene regulatory network inference focus on a particular data set to identify the underlying graph structure, and apply the same method to other data sets and so on. Or, clustering methods are used on various data sets to find subgroups or classify them. However, we would argue that there are deep relationships between clustering and network inference and they can potentially cover each other’s shortcomings. Spatio-temporal gene expression patterns result from both the network structure and the integration of regulatory signals through the network [17]. Moreover, by comparing gene expression levels in the various perturbation conditions, we might reveal the subtype graph structure and understand heterogeneity across various perturbations.

In this paper, we demonstrate that our RPCA-based method helps to find distinct subtypes and classify data sets in a robust way. In order to interpret multi-dimensional spatio-temporal data sets, it is common to compare the responses over experiments and find differences by looking raw data. As the dimension of high-throughput data increases, this is not possible in practice. The proposed method provides a way to interpret multi-dimensional data sets. The low-rank representation provides the large-scale features and the sparse component shows the interesting details such as genomic aberration-specific responses. The intuition behind this is that one can recover the principal components of a data matrix even though a positive fraction of its entries are arbitrarily corrupted or a fraction of the entries are missing as well [9].

Also, although there is a wealth of literature describing canonical cell signaling networks, little is known about exactly how these networks operate in different cancer cells. Therefore, a possible extension of the proposed method is that once we extract common responses, we apply inference algorithms to identify the unified structure using these common responses. Or, we can also focus on individual sparse components to identify the heterogeneity of network structure across cells of different type. Advancing our understanding of how these networks are deregulated across cancer cells and different targeted therapies will ultimately lead to improve effectiveness of pathway-targeted therapies.

## Conclusion

In this study, we develop a new method for clustering and analyzing multi-dimensional biological data. We illustrate how the proposed method can be useful to extract common event-related neural features across many experimental trials. Also, we show that the proposed method helps to find distinct subtypes and classify data sets in a robust way by separating common response and abnormal responses without any prior knowledge. We are currently applying our method to analyze and cluster RPPA data set of the HER2 positive breast cancer and trying to identify underlying graph structures.

## Methods

### Random Projection (RP) and Identifiability

#### Random Projection(RP)

Recent theoretical work has identified random projection as a promising dimensionality reduction technique [19]. Projecting the data onto a random lower-dimensional subspace preserves the similarity of different data vectors, for example, the distances between the points are approximately preserved. Also, RP can reduce the dimension of data while keeping clusters of data points well-separated [19]. Moreover, using RP is substantially less expensive to compute than using techniques such as PCA (Principal Component Analysis) because RP is data-independent.

The idea of RP is that a small number of random linear projections can preserve key information. Theoretical work [19] [20] [21] [22] guarantees that with high probability, all pairwise Euclidean and geodesic distances between points on a low-dimensional manifold are well-preserved under the mapping Ψ : ℝ^*n*^ → ℝ^*m*^, *m* < *n*. Consider a linear signal model

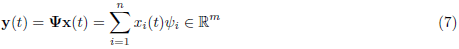

where Ψ = [*ψ*_1_ *ψ*_2_ … *ψ*_*n*_] is an *m* × *n* projection matrix whose elements are drawn randomly from independent identical distributions. First, note that the dimensionality of the data **x** is reduced since *m* < *n*. Also, if we define 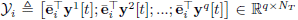 where **ē**_*i*_ is *m*-dimensional unit vector and 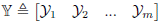, then 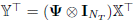 or 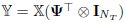 where ⊗ represents the Kronecker product and 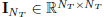 is an identity matrix.

In [19], Dasgupta showed that even if the original distribution of data samples is highly skewed (having an ellipsoidal contour of high eccentricity), its projected counterparts will be more spherical. Since it is conceptually much easier to design algorithms for spherical clusters than ellipsoidal ones, this feature of random projection can simplify the separation into the low-rank and sparse components. Therefore, we can reduce the computational complexity of the non-smooth convex optimization, in particular *l*_1_ and nuclear norms minimization, used in RPCA^1^.

#### Identifiability

Suppose our input 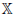 in equation (6) can be decomposed as 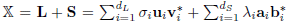 where *σ*_*i*_ are the positive singular values, **u**_*i*_ ∈ ℝ^*q* × 1^, 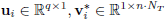 are the left- and right-singular vectors of **L**, and *d*_*L*_ represents the rank of the matrix **L**. *d*_*S*_ is the number of sparse components in **S**, and 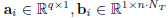 are sparse with only one nonzero entry respectively. By using RP, we have for 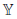,

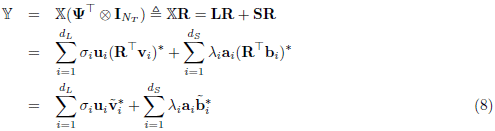

where we denote 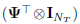 by **R**. As we mentioned above, our input 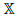 is sparse, so the singular vectors of the low-rank matrix **L** might not be reasonably spread out. However, by using RP (multiplying by **R**), the singular vectors 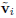 of the resulting matrix become reasonably spread out.

### Cluster Analysis

#### Overview: Dissimilarity

Common measures of dissimilarity for data include Euclidean distance [13], 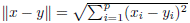 where *x* and *y* are *p*-vectors of measurements on the objects to be clustered. Also, Manhattan distance 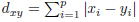 is used, and the “1-*correlation*” distance is defined as follows

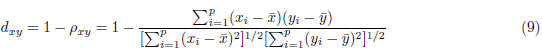

The 1-*correlation* distance is bounded in [0, 2]. This dissimilarity is invariant to changes in location or scale of either *x* or *y*. The 1-*correlation* dissimilarity can be related to the more familiar Euclidean distance: if 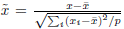 and 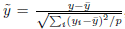, then 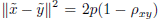. That is, squared Euclidean distance for standardized objects is proportional to the correlation of the original objects. For microarray data, the choice of a dissimilarity measure makes it a popular choice for biological applications. Changes in the average measurement level or range of measurement from one sample to the next are effectively removed by this dissimilarity.

#### Missing data and corruption

As we mentioned, in microarray data, missing data and corrupted data are quite common so in order to deal with missing values, one can modify clustering methods or impute a “complete” data set before clustering. For example, we consider highly-correlated signal *x*_*L*_ = sin(*t*) + *n*_1_ and *y*_*L*_ = sin(*t*) + *n*_2_ where *t* is time step and *n*_1_, *n*_2_ are Gaussian noise 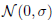. Now, we add a sparse corruption (*x*_*S*_) to the original signal (*x*_*L*_) as shown in Figure 10(a) and calculate the dissimilarity between *x*_*corr*_(= *x*_*L*_ + *x*_*S*_) and *y*_*corr*_ (= *y*_*L*_ + 0). Even though we choose the *d*-sparse corruption of *x*_*S*_ where *d*(≪ *p*) is the number of nonzero component in *x*_*S*_, the correlation is degraded as shown in Figure 10(b) (left). Assuming that we know the corruption signal *x*_*S*_ and *y*_*S*_, we can decompose *x*_*corr*_, *y*_*corr*_ as *ϕ* = [*x*_*L*_; *x*_*S*_] ∈ ℝ^2*p*^ and *ψ* = [*y*_*L*_; *y*_*S*_] ∈ ℝ^2*p*^ respectively. In Figure 10(b)(middle), the red square represents the corruption signal where *y*_*S*_ = 0. Since corruption signal changes the mean and the variance, the correlation is still degraded in (b) (middle). We introduce *γ* so that we allow different weighting factors for (*x*_*L*_, *y*_*L*_) and (*x*_*S*_, *y*_*S*_) respectively. For example, we choose small *γ* for the corruption signal (*x*_*S*_, *y*_*S*_).

**Figure 10.**
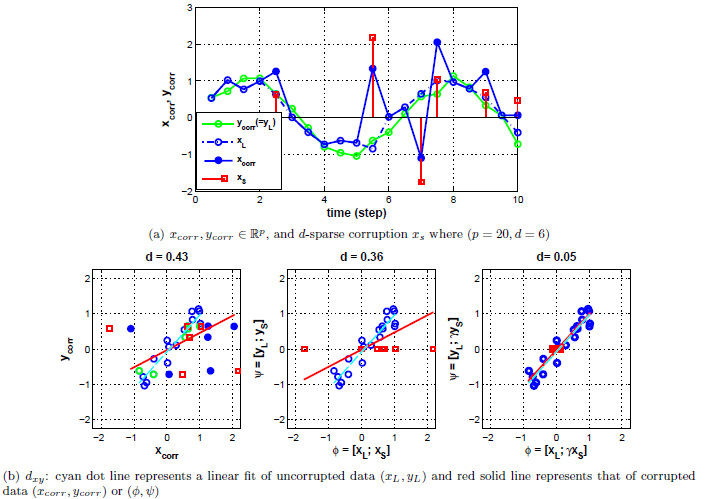
Simple example: (a) green solid line with circle (-○-) represents *y*_*corr*_(= *y*_*L*_ + 0) and blue solid line with circle (-•-) represents *x*_*corr*_(= *x*_*L*_ + *x*_*S*_) where filled circle (•) represents corrupted data, unfilled circle (○) represents uncorrupted data (*x*_*L*_) and unfilled square (□) represents corruption signal (*x*_*S*_) (b) *x*_*corr*_-*y*_*corr*_ plot with 1-*correlation* distance (*d*_*xy*_) without modification(left), with disentanglement(middle), and with disentanglement/weighting factor *γ*.

Therefore, in order to deal with corrupted signals and cluster them, we should separate the original signal and corruption signal first and then calculate the dissimilarity with adjusting weighting factor *γ*. For a gene expression time series data set, when a gene is knocked out, systems are subjected to controlled perturbations and the broad gene expression levels of other genes are perturbed. We can reveal extra information by comparing gene expression levels in the perturbed system with those in the original system. Since abnormal behaviors or different responses to external stimuli or different cell lines can be extracted from the original data using the information available in the data set, we could cluster data and reveal biological meaningful patterns.

#### Our approach: a new *1-correlation* distance

We rewrite the “1-*correlation*” distance (9) as 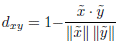 where *x*, *y* ∈ ℝ^*p*^, 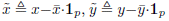 and 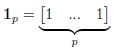 and consider the separation as follows: 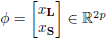 and 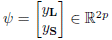 where *x* = *x*_**L**_ + *x*_**S**_, *y* = *y*_**L**_ + *y*_**S**_ and the subscript **L**, **S** represent low-rank component and sparse component. We define “1-*correlation*” distance for *ϕ*, *ψ*, as follows:

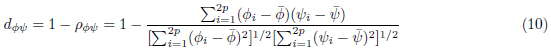

where 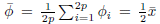 and 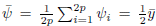. The relation between *d*_*xy*_(= 1 − *ρ*_*xy*_) and *d*_*ϕψ*_(= 1 − *ρ*_*ϕψ*_) is as follows:

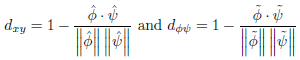

where 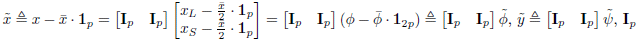 is *p*-dimensional identity matrix, 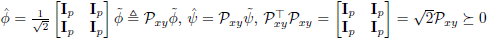 and 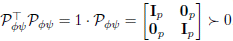.

Therefore, *d*_*xy*_ uses the mixture of low-rank component and sparse component but *d*_*ϕψ*_ calculates the correlation based on the separation. Also, in order to adjust the weighting factor as shown in Figure 10(b) (right), we simply denote 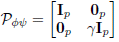 where *γ* is a weighting factor.

**Lemma 1.** *If the sparse component is zero, d*_*ϕψ*_ = *d*_*xy*_.

*Proof*. Since *x*_*S*_ = 0 and *y*_*S*_ = 0, we can simply consider *ϕ*, *ψ* as 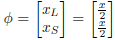 and 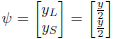 respectively and *γ* = 1

For the disentanglement, we propose the RPCA-based (Robust Principal Component Analysis, [9]) method which uses the information available in the data set in order to identify similar expression patterns^2^.

## Acknowledgments

This research was supported by the NIH NCI under the ICBP and PS-OC programs (5U54CA112970-08), the NIGMS and by the NSF under grant EFRI 1137267.

## Supporting information

**Figure S1.**
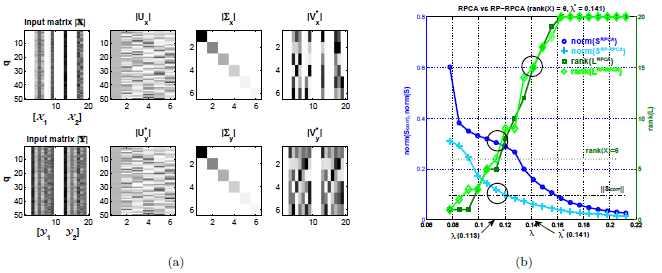
(a) (upper) Input matrix 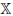 and singular value decomposition (SVD) 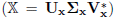. (lower) Randomly projected input matrix 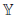 and SVD 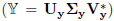. Note that since rank(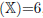, **U**_*x*_ ∈ ℝ^*q* × 6^, **Σ**_*x*_ ∈ ℝ^6 × 6^, 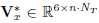. In order to show how well singular vectors are spread out, we show the absolute value of each component. White represents zero value. (b) RPCA results. We run RPCA for sparsely corrupted 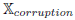, 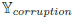. (We added sparse corruption to 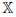 as shown in Figure S2.) Left *y*-axis represents the norm of 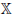 − **L** and the right *y*-axis shows the rank of **L**.

**Figure S2.**
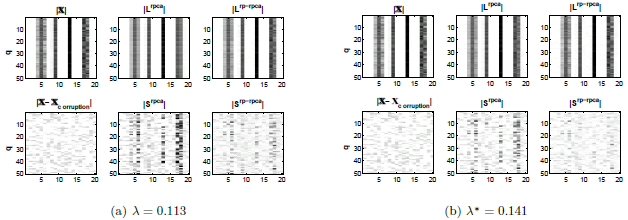
The out of RPCA and RP-RPCA with two different λ values: (a) For λ = 0.113, both **L**^rpca^ and **L**^rp-rpca^ have rank 6 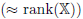 as shown in Figure 4(b). There is a big difference between **S**^rpca^ and the constructed corrupted signal 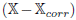 (b) For λ^*^ = 0.141, **S**^rp-rpca^ is close to 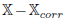 but the low-rank components are misidentified by both RPCA and RP-RPCA because both **L**^rpca^ and **L**^rp-rpca^ have rank 15. Therefore, for RP-RPCA, the separation of the low-rank component and sparse component is close to the true solution but for original RPCA, we have misidentification in both the low-rank and sparse components. We can easily see that **S**^*rpca*^ shows characteristics of the low-rank component in Figure S2 (middle columns of each panel).

**Figure S3.**
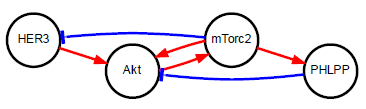
Abstract HER2 overexpressed breast cancer model.

**Figure S4.**
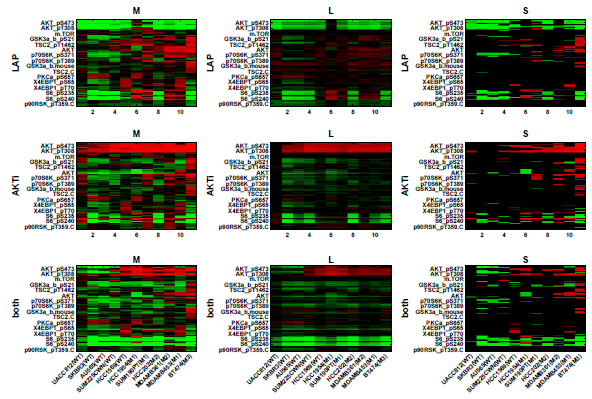
Separation result: (1_*st*_ column) raw data (2_*nd*_ column) low-rank component and (3_*rd*_ column) highly corrupted sparse component using threshold (`M1: H1047R` (kinase domain mutation), `M2: E545K` (helical domain mutation), and `M3: K111N` mutation in `PIK3CA`).

## Footnotes

1 Many speedup methods were developed in optimization by avoiding large-scale SVD. In [23], Mu *et al*. demonstrated the power of projected matrix nuclear norm by reformulating RPCA and in [24], Zhou *et al*. demonstrated the effectiveness and the efficiency of Bilateral Random Projections. However, both methods consider a dense matrix 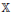 while in this paper we consider the case when the input matrix 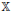 is sparse.

2 In [18], Liu *et al*. proposed an RPCA-based method of discovering differentially expressed genes using static data. They provided an efficient and effective approach for gene identification. However, we focus on the spatio-temporal gene expression data set and consider the disentanglement of low-rank and sparse component to extract common features and detect specific response or heterogeneity via modified RPCA. Here, we treat the spatio-temporal gene expression and focus on the relationship between gene regulatory network and dynamics of regulatory signal. We note this goes beyond the results in [14] due to the transformation involved.

